# pTRIP, a novel integration plasmid for *Listeria monocytogenes*

**DOI:** 10.64898/2026.09.11.750939

**Authors:** Jessica Schüler, Jeanine Rismondo

## Abstract

In the past decades, several tools to genetically modify the human pathogen *Listeria monocytogenes* were developed. Here, we constructed a new integrative plasmid system for *L. monocytogenes* named pTRIP, for *<u>tr</u>eB* insertion <u>p</u>lasmid. pTRIP is a vector which stably integrates into the *treB* locus of the wild type EGD-e. This locus encodes the sole trehalose-specific EIIB and EIIC component of a phosphotransferase system. Successful integration leads to the disruption of *treB* and thus, to an inability of the resulting *L. monocytogenes* strains to grow on trehalose as sole carbon source. Due to integration through double homologous recombination, it is the first integrative system which does not require antibiotic selection pressure. To assess functionality of the pTRIP system, *prfA* and its native promoter region were integrated into the *treB* locus of a Δ*prfA* strain. Complementation was confirmed in 78% of the isolated clones, indicating successful integration of *prfA* into the *treB* locus. We further constructed derivatives of pTRIP harboring the constitutive *P_p60_* (pTRIP1) and the inducible *P_rha_* (pTRIP2) promoter to further expand application possibilities. Microscopic analyses confirmed the functionality of both promoter constructs and showed dose-dependent induction for *P_rha_.* pTRIP is an efficient tool for stable gene expression as well as functional studies and expands genetic modification possibilities for *L. monocytogenes*.

**Importance:** *L. monocytogenes* is an important human pathogen that can cause severe infections in at-risk group individuals such as elderlies or immunocompromised patients. Over the past decades, several genetic tools have been developed to study the physiology and pathogenicity of *L. monocytogenes*. Currently, the possibilities to genetically modify *L. monocytogenes* using integrative plasmids are limited as these plasmids use the same integration site. We constructed the integrative plasmid pTRIP that integrates into the *treB* locus via homologous recombination. This enables stable integration without the use of antibiotic selection pressure leaving flexibility for usage of additional plasmids such as pPL2 or pIMK3. The pTRIP system therefore expands the genetic toolbox for *L. monocytogenes* and will be useful for future studies.

## Introduction

*Listeria monocytogenes* is a Gram-positive, rod-shaped bacterium. It can live as a saprophyte, is ubiquitous in nature and is able to inhabit various environments. Upon ingestion of contaminated foods, the bacterium can enter hosts such as humans and ruminants, where it expresses its virulence factors and establishes infection. In its pathogenic lifestyle, *L. monocytogenes* can cause a self-limiting gastroenteritis in healthy individuals but is a threat especially to susceptible person groups such as immunosuppressed, old or pregnant people. In this case, infections can lead to severe listeriosis manifesting in sepsis, meningitis, encephalitis and abortions. Listeriosis reaches a hospitality rate of over 90% with a 20-30% mortality rate which establishes it as a particularly serious form of foodborne disease (1–3). The pathogen has been studied for almost a century and is a well-established model organism regarding bacterial pathogenesis and pathophysiology (4, 5). The availability of genetic tools including integrative and allelic exchange vectors have highly contributed to the characterization of *L. monocytogenes* and have facilitated investigations of its physiology, virulence and host-pathogen interactions (4). Known integrative vectors, such as pPL2 or pIMK plasmids are limited to one integration site in EGD-e and 10403S (6, 7), which are the main wild type strains of *L. monocytogenes* used worldwide (8). These plasmids are ectopically integrated into the *attP* site of the *tRNA^Arg^* locus using the PSA phage integrase (6, 7). Although these tools provide valuable application, only one integrative plasmid can be used for a single strain. Therefore, we addressed this restriction and aimed to broaden the genetic toolbox for *L. monocytogenes* EGD-e.

In this study, we describe the development of a novel 9,675 bp integration plasmid named pTRIP, which stably integrates into the *treB* locus of *L. monocytogenes* through double homologous recombination without the need for antibiotic selection. The integration of pTRIP leads to disruption of the *treB* gene, which encodes a trehalose-specific EIIB and EIIC component of a phosphotransferase system (9). *L. monocytogenes* strains lacking TreB are unable to import trehalose (9, 10), thus, integration of pTRIP can be confirmed by an inability of the obtained strains to grow on trehalose as sole carbon source. Moreover, we further constructed variants of pTRIP, which enable inducible expression as well as constitutive expression of the gene of interest. By providing an additional chromosomal integration site, pTRIP and its derivatives expand the genetic toolbox for the *L. monocytogenes* wild type EGD-e and enable the construction of strains carrying pTRIP in combination with other integrative plasmids, thereby supporting more complex genetic modifications.

## Results

### A new *treB* targeting integration plasmid (pTRIP) expands insertional cloning in EGD-e

So far, all known insertion plasmids integrate into the *tRNA^Arg^*locus which allows only a single integration event under constant selection pressure. To overcome this, a novel 9,675 bp integration plasmid was constructed in this study named pTRIP for ***<u>tr</u>****eB* **i**nsertion **<u>p</u>**lasmid. This plasmid expands the toolbox to genetically modify the *L. monocytogenes* wild type EGD-e. It is based on the backbone of the suicide plasmid pHoss1, which contains an antisense *secY* RNA expression cassette facilitating screening for colonies that lost the plasmid (11, 12). Moreover, pTRIP contains the disrupted *treB* locus from EGD-e split by the multiple cloning site (MCS) from pPL2 (6, 9, 10) (Figure 1 A). The plasmid stably integrates into the *treB* locus (*lmo1255*), which encodes the sole trehalose PTS permease (9), via double homologous recombination. Deletion of *treB* was shown to result in an inability of *L. monocytogenes* to grow on trehalose as sole carbon source (10). Thus, successful integration of pTRIP can be verified by screening on trehalose-containing minimal medium, providing a simple and efficient method for selecting correctly integrated clones. In comparison to the commonly used integration vectors such as pPL2 and pIMK2 (6, 7), pTRIP facilitates a stable chromosomal integration of the gene of interest without the need to constantly maintain the plasmid inside the cell.

**Fig. 1:**
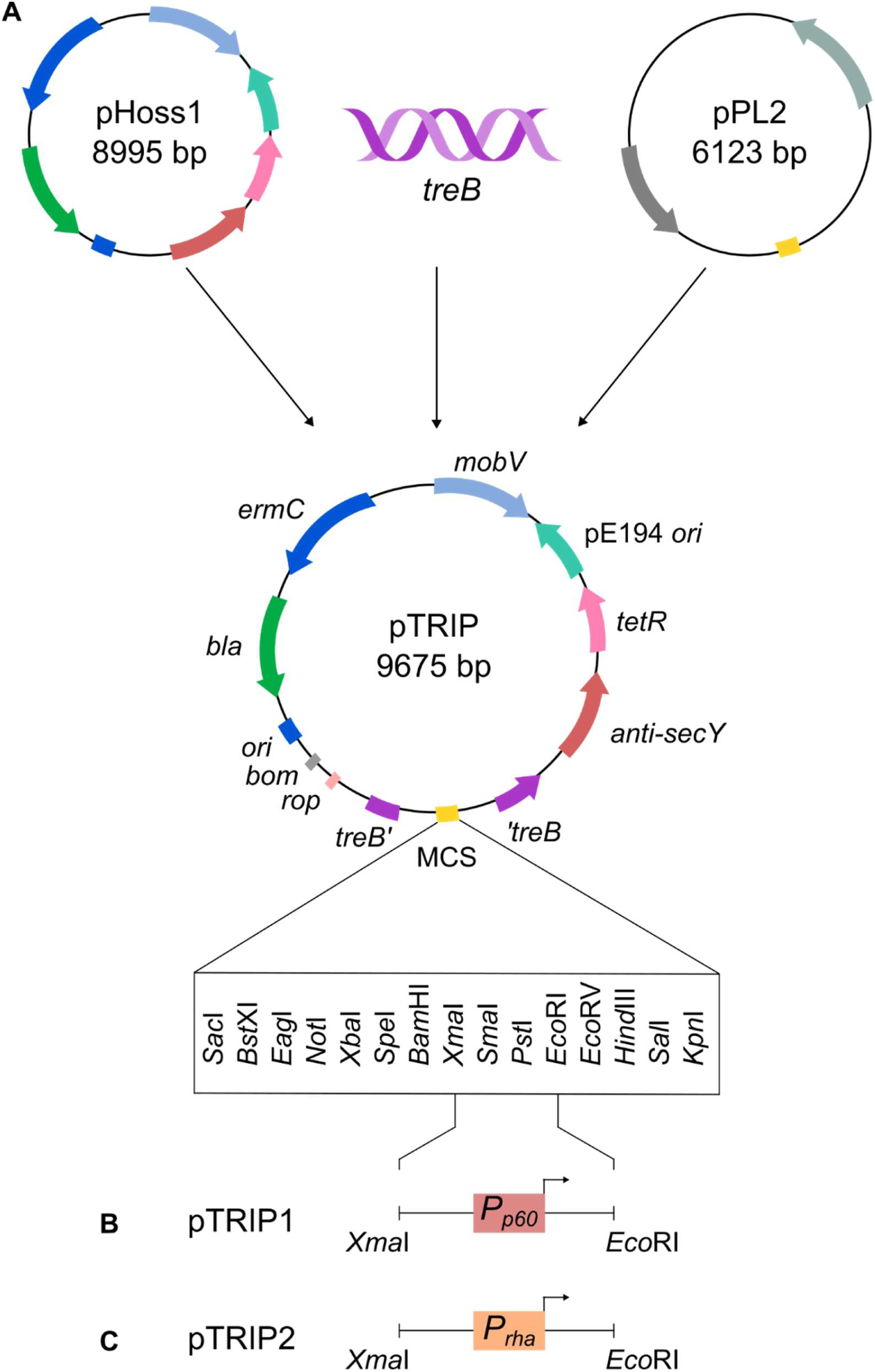
Plasmid map of the new *treB* integration plasmid pTRIP. (A) The novel plasmid is derived from the backbone of the suicide plasmid pHoss1, a disrupted *treB* locus from EGD-e and the multiple cloning site (MCS) from pPL2. Unique restriction sites of the MCS are shown in a box below the plasmid. (B) pTRIP1 with a constitutive *P_p60_* promoter amplified from pIMK2 and cloned into the MCS with *Xma*I and *Eco*RI. (B) pTRIP2 with an inducible *P_rha_* promoter amplified from EGD-e and cloned into the MCS via *Xma*I and *Eco*RI. *anti-secY*: tetracycline-inducible antisense Y gene; *tetR*: tetracycline repressor; pE194 *ori*: pE194 replication origin; *mobV*: mobilization gene V; *ermC*: erythromycin resistance cassette; *bla*: ampicillin resistance cassette; *ori*: origin of replication; *bom*: basis of mobility region; *rop*: repressor of primer.

To further expand the usability of the pTRIP system, different promoters were inserted into the MCS via *Xba*I and *Eco*RI restriction sites. Using this approach, multiple plasmid variants were constructed: pTRIP without a promoter to allow expression under the native promoter of the gene of interest, pTRIP1 containing the constitutive promoter *P_p60_* and pTRIP2 containing the rhamnose-inducible promoter *P_rha_* (7, 13)(Figure 1 B+C).

### Integration into the *treB* locus does not affect biofilm formation and acid survival in EGD-e

Previous studies have shown that inactivation of TreB by an N352K substitution not only resulted in an inability to metabolize trehalose, it also affected biofilm development and acid resistance in strain 1386, a clonal complex 5 (CC5) strain (9). To ensure that *treB* is a suitable insertion locus for *L. monocytogenes* EGD-e, an insertion strain was constructed where the MCS of the pTRIP vector was inserted into the *treB* locus yielding strain *treB*::MCS. Its growth and its ability to survive acid stress and form biofilms was compared to the *L. monocytogenes* wild type strain EGD-e and a *treB* deletion strain.

Growth of the *treB*::MCS strain and the Δ*treB* strain was similar to the wild type in BHI, and LSM with glucose or cellobiose as sole carbon source, respectively (Figure 2A-C). As expected, disruption of *treB*, the sole trehalose PTS permease in *L. monocytogenes* (9, 10), led to a severe growth defect of both Δ*treB* and *treB*::MCS when grown in LSM supplemented with trehalose as sole carbon source (Figure 2D). Both strains started to grow under these conditions after approximately 22 h, indicating the formation of suppressors.

**Fig. 2:**
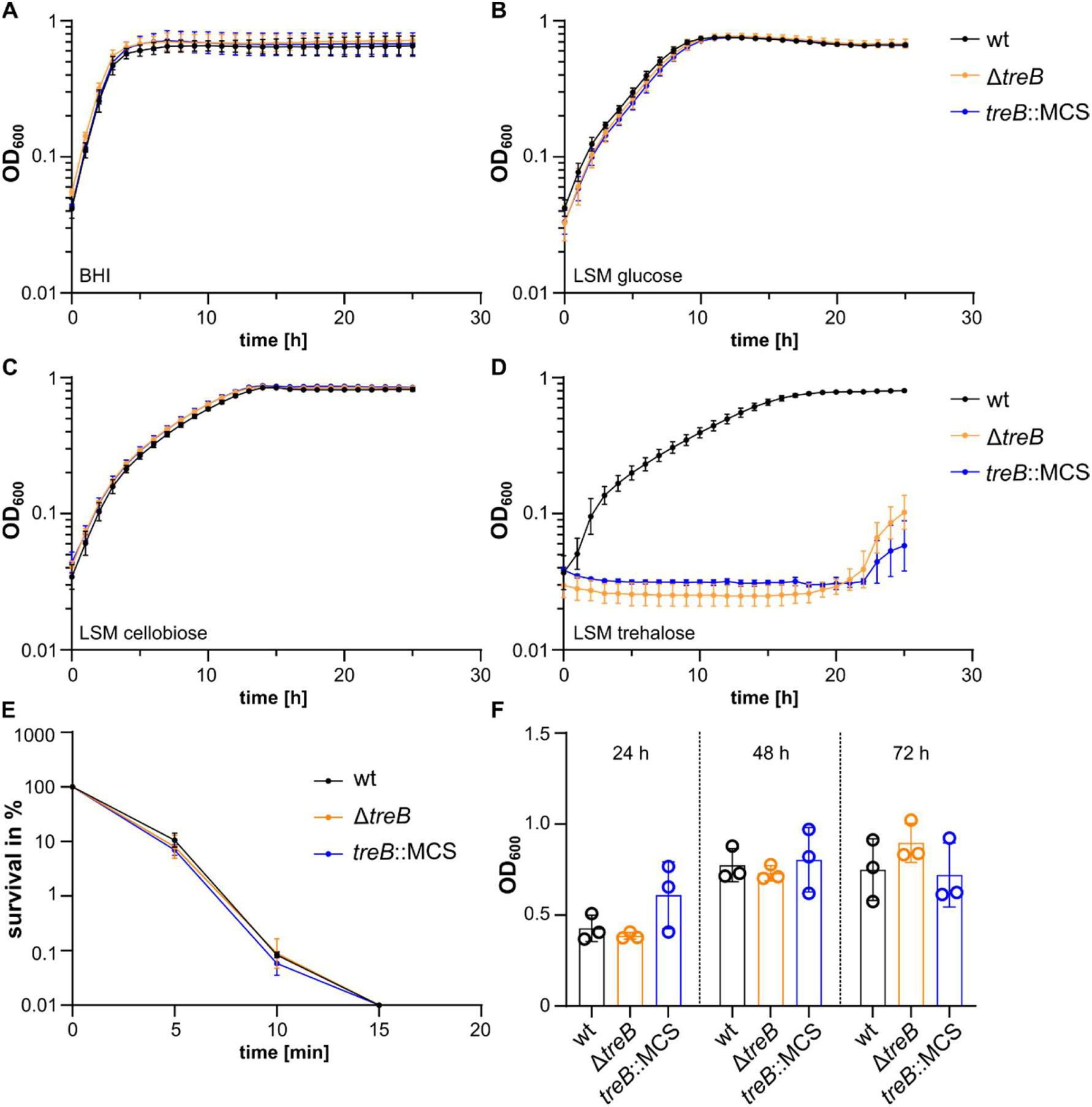
Physiological effects of *treB* disruption. The indicated *L. monocytogenes* strains were grown in (A) BHI, (B) LSM supplemented with 1 % glucose, (C) LSM supplemented with 1 % cellobiose and (D) LSM supplemented with 1 % trehalose at 37°C as described in the method section. (E) Acid survival in % after 5 min, 10 min and 15 min in acidified BHI (pH 2.3). (F) Biofilm production after 24 h, 48 h and 72 h at 30°C. For growth and acid survival assays, the averages and standard deviations of three biological replicates are shown. For the biofilm assay, averages of six technical replicates are shown for biological triplicates, respectively, with their standard deviation.

A lack of *treB* has been shown to decrease acid resistance in strain 1386 which indicated that trehalose metabolism might affect acid survival of *L. monocytogenes* (9). An acid survival assay was conducted and strains were exposed to acidified BHI (pH 2.3) for 5, 10 and 15 minutes to observe survival under acid stress. Interestingly, sensitivity to acid in EGD-e was not affected by *treB* disruption as *treB*::MCS and Δ*treB* survival was identical to the wild type (Figure 2E). In all strains, survival drastically dropped to 0.1 % after 10 minutes of acid exposure and after 15 minutes exposure, no bacterial colonies could be recovered (Figure 2E).

Disruption of trehalose metabolism was also shown to decrease biofilm production at 30°C in *L. monocytogenes* 1386 (9). To test this for EGD-e, biofilm formation was analysed at 30°C after 24, 48 and 72 h. Biofilm production of *treB*::MCS and Δ*treB* was comparable to that of the wild type (Figure 2F), indicating that trehalose metabolism has no impact on biofilm formation in EGD-e under the tested conditions.

Together, these results demonstrate that insertion of the MCS cassette into the *treB* locus did not affect growth on glucose and cellobiose, acid survival, or biofilm formation. Therefore, *treB* can indeed be used as an insertion site for the pTRIP system.

### Complementation of *prfA* at the *treB* locus

To assess the functionality of pTRIP, pTRIP*-P_plcA_-P_prfA_-prfA* was introduced into a Δ*prfA* strain resulting in the construction of a *prfA* complementation strain (*prfA* compl.). Complementation of *prfA* offers an easy screening method as LB agar plates supplemented with 0.2% activated charcoal and egg yolk serve as a visual indicator of PrfA activity. The activated charcoal in the LB agar activates PrfA, which is normally inactive under these conditions, through an unknown mechanism (14). The expression of *plcB*, which is a virulence gene of *L. monocytogenes*, is directly regulated by PrfA. *plcB* encodes a phospholipase C, which also has a lecithinase activity and is therefore able to hydrolyse lecithin of egg yolk resulting in halo formation. Therefore, PlcB activity serves as a qualitative readout of PrfA function. We previously used the same approach to complement the Δ*prfA* strain with *P_plcA_-P_prfA_-prfA* using pPL3e, one of the commonly used vectors that integrates into the *tRNA^Arg^* locus (10), thus, serving as an ideal strategy to assess the functionality of pTRIP. To analyse the rate of successful integration, the integration protocol was performed using three biological and four technical replicates, yielding a total of 559 tested strains. For all tested pTRIP derivatives, the integration protocol resulted in a 100% plasmid loss rate upon induction of *secY* expression with 1.5 µg/ml anhydrotetracycline (AHT) (data not shown). Successful re-integration of *prfA* was achieved with an efficiency of 78%. Screening for successful integration was performed using LSM supplemented with trehalose as sole carbon source and additionally LB agar supplemented with 0.2% activated charcoal and egg yolk to screen for restored PrfA activity (Fig. 3A). Clones, which were able to grow on LSM trehalose, did not show a halo on LB agar supplemented with activated charcoal and egg yolk indicating PrfA inactivity and thus unsuccessful pTRIP*-P_plcA_-P_prfA_-prfA* integration. It should be noted that some of the patched colonies with restored PrfA activity were still able to show slight growth on LSM trehalose, likely because they can use nutrients of dead cell material. Therefore, only strong growth on trehalose should be considered as an indicator of unsuccessful plasmid integration (Fig. 3A, stars). To further verify wild type-like PrfA activity for the *prfA* complementation strain, a PlcB test was performed, where PrfA activity was also assessed in the presence of PTS-dependent sugars. PTS-dependent sugars such as glucose or cellobiose are known to inhibit PrfA activity through the phenomenon of sugar-dependent PrfA inhibition (2, 15). As expected, the *prfA* complementation strain shows halo formation on LB charcoal plates without sugar addition but not when glucose or cellobiose were added and thus behaves identically to the wild type EGD-e (Figure 3B). Thereby, successful complementation was confirmed and the functionality of the pTRIP system demonstrated.

**Fig. 3:**
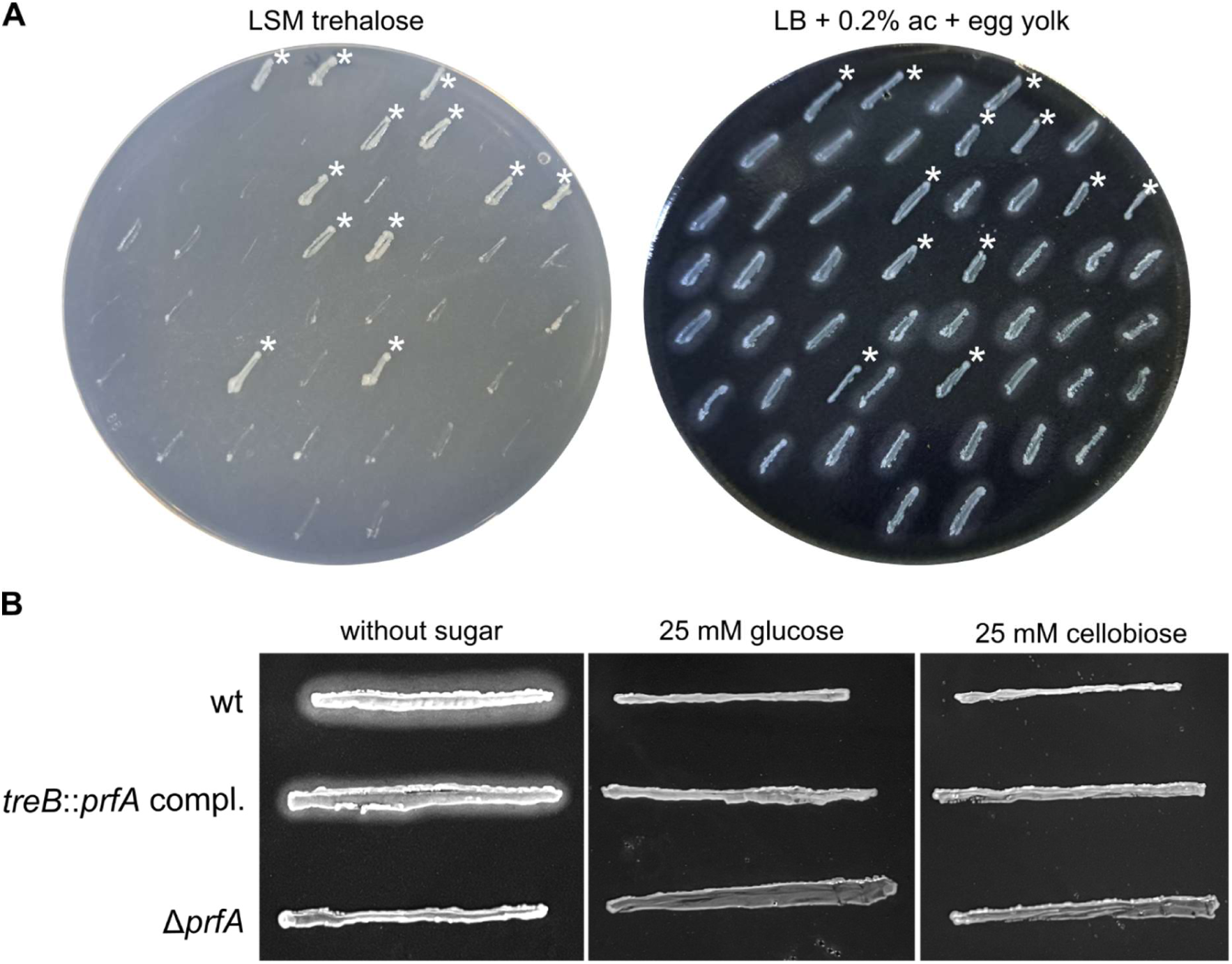
Complementation of Δ*prfA* using pTRIP. (A) Patched clones on LSM supplemented with 1% trehalose and on LB agar supplemented with 0.2% activated charcoal (ac) and egg yolk. Colonies which showed strong growth on LSM trehalose and without halo on LB with activated charcoal and egg yolk are marked with a star (*) and represent unsuccessful integration. A representative image of three biological replicates with four technical replicates each is shown. (B) PlcB activity of the EGD-e *treB::prfA* complementation strain, EGD-e and Δ*prfA*. The strains were streaked on LB agar supplemented with 0.2% activated charcoal, egg yolk and either 25 mM glucose or cellobiose, or no added sugar and incubated at 37°C. A representative image of three biological replicates is shown.

### Promoter constructs enable constitutive and inducible gene expression

To measure promoter activity, the *msfGFP* gene was cloned downstream of the respective promoters and transformed into the wild type EGD-e. For the *P_rha_* promoter (pTRIP2), rhamnose was added in increasing concentrations to observe dose-dependent induction of fluorescence. The rhamnose operon is subject to carbon catabolite repression (CCR), therefore cultures can only be cultivated in media lacking glucose, such as LB medium (13). As a control, EGD-e was transformed with pTRIP-*msfGFP* lacking a promoter which, as expected, did not show fluorescence. Increasing rhamnose concentrations led to a gradually increase in fluorescence with a concentration of 0.05% rhamnose being sufficient to induce weak promoter activity of *P_rha_* in a subpopulation of cells (Figure 4). At 0.5% rhamnose, an increase of fluorescence was observed and led to a fluorescent signal in all cells. As expected, no fluorescence signals can be observed when rhamnose and glucose were added due to CCR. The constitutive promoter *P_p60_* in pTRIP1-*msfGFP* resulted in constitutive expression of *msfGFP* and led to fluorescence signal in all cells. pTRIP1 thus enables constitutive overexpression and can be used independently of the medium.

**Fig. 4:**
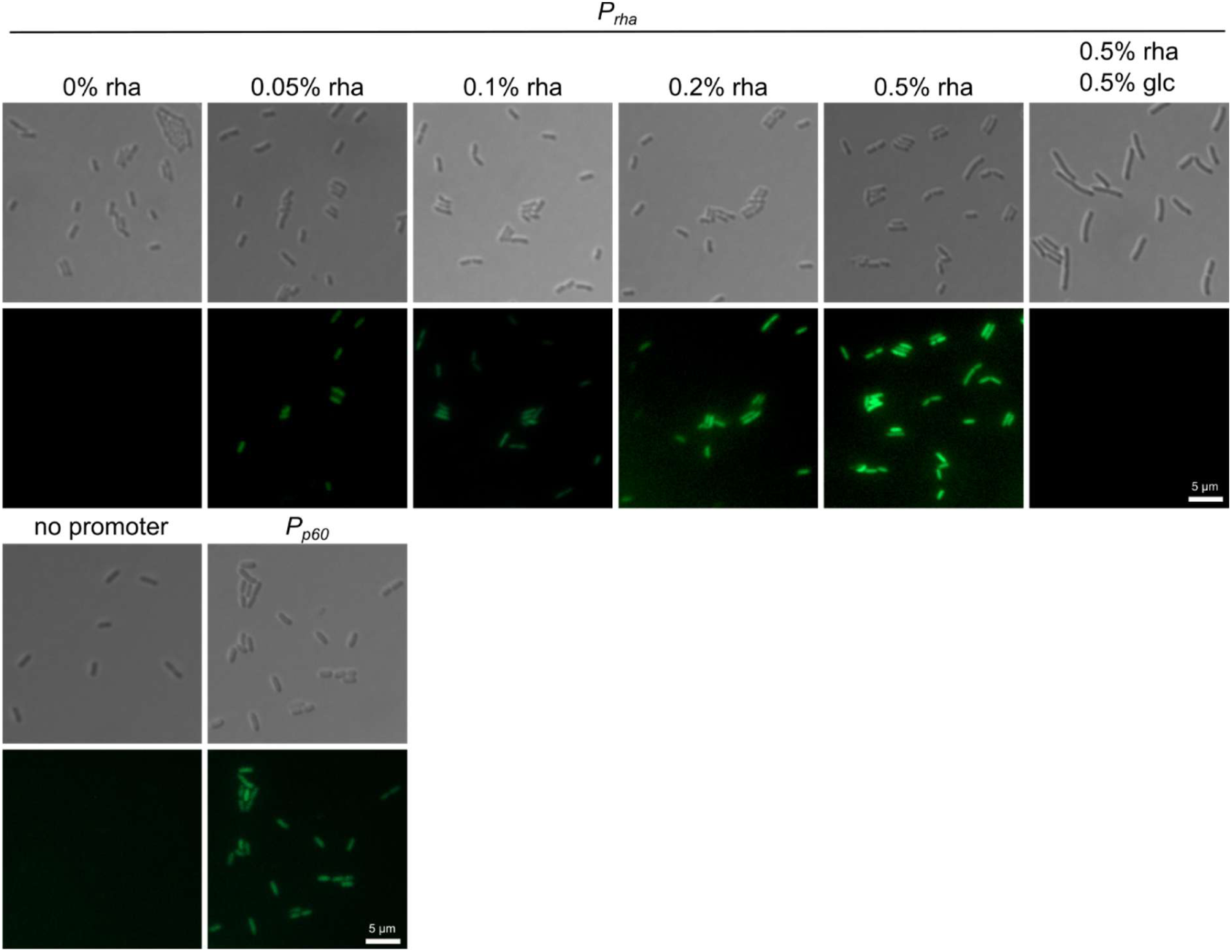
Rhamnose-inducible and constitutive *msfGFP* expression. *P_rha_* activity was monitored through dose-dependent *msfGFP* induction and *P_p60_* activity through constitutive *msfGFP* expression in EGD-e after 20 h of incubation at 100-fold magnification. Rhamnose concentrations are indicated. The top row shows differential interference contrast (DIC) microscopy while the bottom row represents fluorescence microscopy. pTRIP without promoter served as control. rha: rhamnose; glc: glucose. Representative images of three biological replicates are shown.

## Discussion

Integrative cloning approaches have strongly contributed to the characterization of *L. monocytogenes*. However, they are often limited as known integrative vectors such as pIMK or pPL2 plasmids target the *tRNA^Arg^* locus and rely on constant selection pressure which allows only a single integration at a time (6, 7). To address this, a new integrative vector called pTRIP (***<u>tr</u>****eB* **i**nsertion **<u>p</u>**lasmid) was constructed in this study to expand the toolset to genetically modify the *L. monocytogenes* wild type EGD-e.

The pTRIP system provides a useful tool for future studies by enabling chromosomal integration without the need for continuous selection pressure due to stable double homologous integration into the *treB* locus. In addition, cloning with pTRIP has a high success rate (78%) and it is easy to screen for positive clones due to the suicide system from pHoss1 (12)(100% plasmid loss rate) and the inability of desired strains to grow on trehalose as sole carbon source. While the integration of genes of interest with the pTRIP plasmids is more time-consuming than with established integration plasmids, this system greatly expands the possibilities for the genetic modification of *L. monocytogenes,* especially since the resulting strains do not contain an antibiotic resistance cassette. This enables the simultaneous use of other plasmids for integration or gene deletion. Compared to existing integration plasmids, pTRIP is comparatively large (9,675 bp). Therefore, it might not be suitable for the integration of large genes or operons.

Wu et al observed that deletion of *treB* affected several physiological characteristics such as biofilm formation and acid resistance in the CC5 strain 1386 (9). We demonstrated that application of the pTRIP vector does not affect biofilm formation, acid survival and growth on glucose or cellobiose in the wild type EGD-e. The discrepancies between our results and those reported by Wu et al. for the *L. monocytogenes* isolate 1386 might be explained by the genetic background of the strains: EGD-e belongs to lineage II, clonal complex 9 and serotype 1/2a, whereas strain 1386 belongs to lineage 1, clonal complex 5 and serotype 4b (8, 16).

Derivatives of pTRIP enable constitutive and inducible expression of the gene of interest. Fluorescent microscopy analyses confirmed the functionality of the *P_p60_* promoter (pTRIP1) and the inducible *P_rha_* promoter (pTRIP2). It should be noted that *P_rha_* is subject to CCR (13), therefore, this system is not suitable for investigating mechanisms which require glucose and media usage is limited.

Taken together, targeting the *treB* locus with pTRIP is a suitable tool for future studies as it does not seem to exert downstream effects and expands the possibilities for genetic modifications in *L. monocytogenes*.

## Materials and methods

### Bacterial strains and growth conditions

All strains and plasmids used in this study are listed in Table 1. *Escherichia coli* strains were grown in lysogeny broth (LB) medium and *Listeria monocytogenes* strains in brain heart infusion (BHI) medium at 37°C unless otherwise stated. When required, media were supplemented with antibiotics at the following concentrations: For *E. coli* cultures: 100 µg/ml ampicillin (amp); for *L. monocytogenes* cultures: 7.5 µg/ml chloramphenicol (cat), 5 µg/ml erythromycin (erm), 1.5 µg/ml anhydrotetracycline (AHT).

**Table 1:** Bacterial strains used in this study.

| Unique ID | Strain name and resistance | Source |
| --- | --- | --- |
| ***Escherichia coli* strains** |  |  |
| ANG124 | DH5α pKSV7; ampR | (19) |
| ANG1274 | XL1 Blue pPL2; catR | (6) |
| ANG4970 | XL1 Blue pHJS105; ampR | (20) |
| EJR216 | DH5α pHoss1; ampR | (12) |
| EJR274 | XL1 Blue pPL3e-*P<sub>plcA</sub>*-*P<sub>prfA</sub>*-*prfA*; catR | (10) |
| EJR389 | DH5α pKSV7-*ΔtreB*; ampR | (10) |
| EJR405 | DH5α pTRIP; ampR | This study |
| EJR418 | DH5α pTRIP-*P<sub>plcA</sub>*-*P<sub>prfA</sub>*-*prfA*; ampR | This study |
| EJR457 | DH5α pTRIP-*P<sub>rha</sub>*; ampR | This study |
| EJR458 | DH5α pTRIP-*P<sub>p60</sub>*; ampR | This study |
| EJR462 | DH5α pTRIP-*P<sub>rha</sub>*-*msfGFP*; ampR | This study |
| EJR463 | DH5α pTRIP-*P<sub>p60</sub>*-*msfGFP*; ampR | This study |
| EJR464 | DH5α pTRIP-*msfGFP*; ampR (w/o promoter) | This study |
| ***Listeria monocytogenes* strains** |  |  |
| ANG873 | EGD-e | (21) |
| BUG2214 | EGD-e *ΔprfA* | (22) |
| LJR718 | EGD-e *ΔtreB* | (10) |
| LJR749 | EGD-e *treB*::MCS | This study |
| LJR785 | EGD-e $\Delta prfA$ $treB::P_{plcA}-P_{prfA}-prfA$ ($prfA$ compl.) | This study |
| LJR843 | EGD-e $treB::P_{p60}-msfGFP$ | This study |
| LJR845 | EGD-e $treB::P_{rha}-msfGFP$ | This study |
| LJR846 | EGD-e $treB::msfGFP$ (w/o promoter) | This study |

### Construction of pTRIP and its derivatives

All primers used in this study are listed in Table 2. The new 9,675 bp *treB* integration plasmid is a high-copy plasmid which was assembled from five independent DNA fragments. Two fragments of the pHoss1 plasmid (12) were amplified by PCR using primer pairs JS17/18 and JS19/20, respectively, from a pHoss1 backbone which had been digested with *Sal*I and *Eco*RI. The first and second half of *treB* were PCR amplified from genomic DNA of EGD-e using primers JS11/12 and JS13/14, respectively. A 327 bp DNA fragment containing the multiple cloning site (MCS) was amplified with primers JS15 and JS16 from pPL2 which had been linearized with *Bgl*II. All PCR products were purified and assembled using Gibson Assembly®. For the assembly reaction, 2 µl of each PCR product were combined with 10 µl of the Invitrogen™ Gibson Assembly® EX Master Mix (Thermo Fisher) and incubated for 1 h at 50°C. The reaction mix was then transformed into *E. coli* DH5α yielding strain EJR405. Plasmids were first checked by test digestion with *Bam*HI and finally confirmed by whole plasmid sequencing (Microsynth, Göttingen, Germany). To test functionality of pTRIP, *prfA* including its promoters (*P_plcA_-P_prfA_-prfA*) was amplified by PCR using primer pair JR312/257 from pPL3e-*P_plcA_-P_prfA_-prfA* (derived from EJR274) which had been cut with *Nco*I. The resulting PCR product was digested with *Kpn*I and *Sal*I and ligated into pTRIP which had been cut with the same enzymes. Recovery was performed in *E. coli* DH5α yielding strain EJR418. For the promoter construction, the *P_p60_* promoter was amplified from pIMK2 which had been linearized with *Kpn*I, using primers JS41/42. The *P_rha_* fragment (13) was amplified from genomic DNA of EGD-e with primer pair JS46/47. The resulting PCR products were digested with *Xma*I and *EcoR*I and ligated into the pTRIP plasmid that had been cut with the same enzymes. Plasmids pTRIP-*P_p60_* (pTRIP1) and pTRIP-*P_rha_* (pTRIP2) were recovered in *E. coli* DH5α yielding strains EJR458 and EJR457, respectively. To verify functional gene expression from the promoter constructs, *msfGFP* was cloned into pTRIP1 and pTRIP2. pTRIP-*msfGFP* lacking a promoter was constructed as negative control. For this purpose, *msfGFP* was amplified from pHJS105 which had been linearized with *Eco*RI using primer pair JS48/49. The resulting PCR product was subsequently digested with *Kpn*I and *EcoR*I and ligated into pTRIP, pTRIP1 and pTRIP2, which had been digested using the same enzymes, yielding strains EJR464, EJR463 and EJR462, respectively.

**Table 2:**
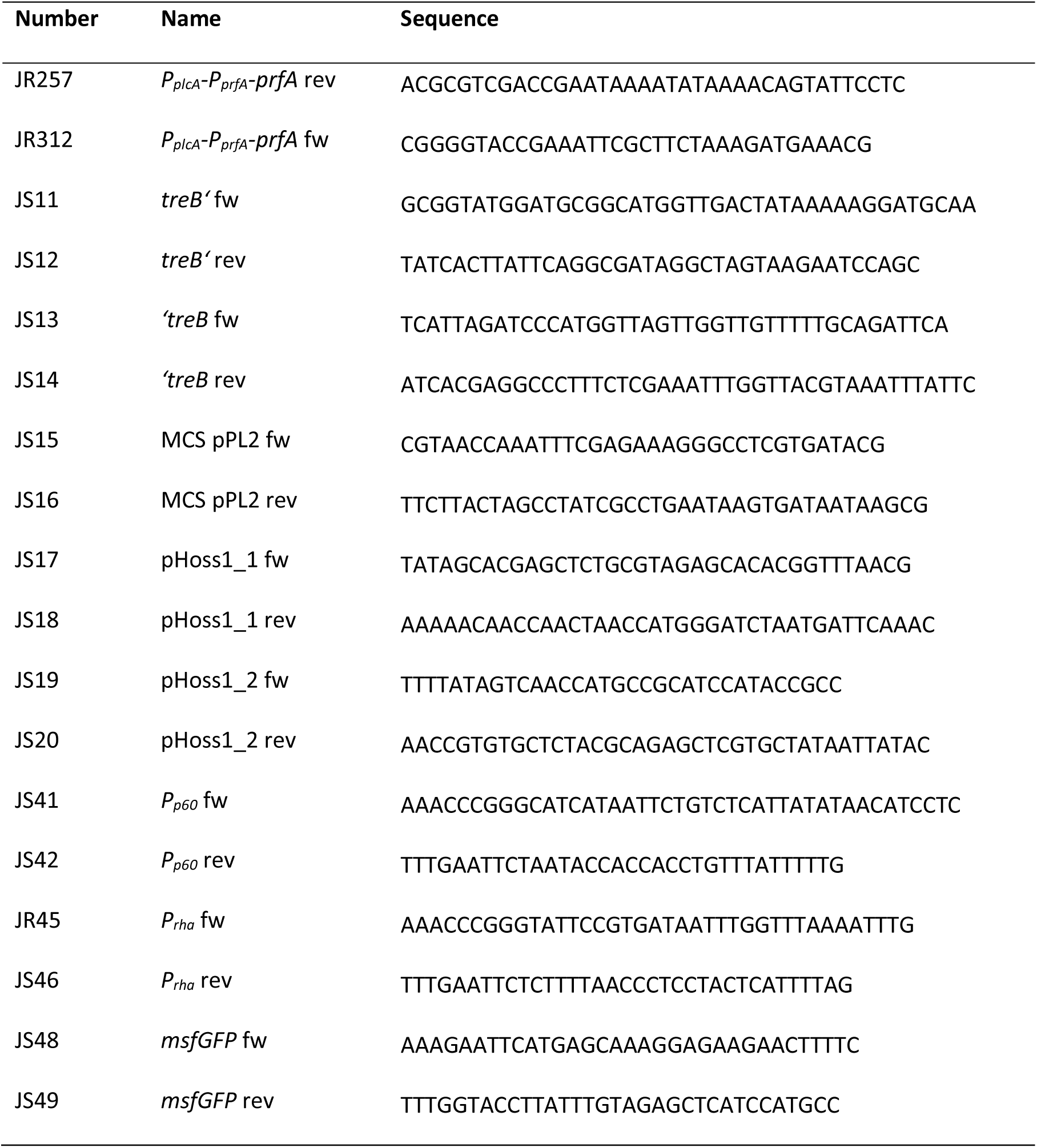
Primers used in this study.

#### Gene insertion with pTRIP

For the construction of *L. monocytogenes* strains carrying genes of interest integrated into the *treB* locus, bacteria were transformed using the novel integration plasmid pTRIP. The integration protocol was adapted from Abdelhamed et al. (12). After transformation, cells were streaked onto BHI agar supplemented with 5 µg/ml erythromycin and incubated for two days at 30°C. Single colonies were then plated on BHI agar supplemented with 5 µg/ml erythromycin and incubated for two days at 42°C. Colonies were repeatedly restreaked on BHI agar supplemented with 5 µg/ml erythromycin and incubated for further two days at 42°C. Afterwards, colonies were inoculated in liquid BHI without antibiotics overnight at 30°C with agitation, a step which was repeated twice. A 1:500 dilution of the overnight culture was inoculated in 4 ml BHI broth and incubated for eight hours at 42°C with agitation. To select for colonies without plasmid, 10^-3^ and 10^-4^ dilutions of the day culture were plated on BHI agar containing 1.5 µg/ml anhydrotetracycline followed by incubation at 30°C for 3 days. From each transformation, 50 colonies were picked and patched onto BHI agar with or without 5 µg/ml erythromycin and LSM agar containing 1% trehalose as sole carbon source (17, 18). Plates were incubated at 37°C overnight. Colonies which only grew on BHI agar were further analysed for successful integration via colony PCR. In order to check for *prfA* complementation, colonies were additionally patched on LB agar supplemented with 0.2% activated charcoal and 1% egg yolk.

To assess the functionality of the integration protocol, pTRIP was transformed into EGD-e yielding strain LJR749. To further verify that pTRIP can be used for the construction of complementation strains, where the corresponding gene is under the control of the native promoter, a Δ*prfA* complementation strain was constructed. For this, pTRIP-*P_plcA_-P_prfA_-prfA* (EJR418) was transformed into *L. monocytogenes* EGD-e Δ*prfA* (BUG2214) resulting in strain LJR775. For microscopic assessment of promoter functionality, plasmids pTRIP-*msfGFP*, pTRIP1-*msfGFP* and pTRIP2-*msfGFP* were transformed into EGD-e resulting in the construction of EGD-e *treB*::*P_p60_-msfGFP* (LJR843), EGD-e *treB*::*P_rha_-msfGFP* (LJR845) and EGD-e *treB*::*msfGFP* (no promoter) (LJR846).

#### Lecithinase (PlcB) assay

PlcB activity was assessed by qualitatively detecting lecithin hydrolysis as an indicator for PrfA activity. Bacterial strains were either patched on LB agar containing 0.2% activated charcoal, 2% egg yolk of an egg yolk suspension, which was prepared by mixing one egg yolk with an equal volume of 1x phosphate-buffered saline (PBS, pH 7.4), and, where indicated, 25 mM glucose or cellobiose. Plates were incubated for 24 h at 37°C and photographed.

### Acid survival assay

To assess bacterial survival under acidic conditions, a survival assay was performed according to Wu et al. with minor modifications (9). Briefly, 100 µl of an overnight culture were added to 900 µl BHI medium acidified to pH 2.3 with 1 M HCl. Samples were incubated statically at 37°C for 5, 10 and 15 min. At each time point, cultures were serially diluted (10^-1^-10^-7^) in 1x PBS. 10 µl of the undiluted sample and each dilution were spotted onto BHI plates and incubated overnight at 37°C. Survival was calculated as the percentage of the initial cell count. Biological triplicates each with three technical replicates were performed.

### Biofilm assay

Biofilm assays were conducted as previously described with minor modifications (9). Overnight cultures were adjusted to an OD_600_ of 0.01 in BHI broth. 200 µl were transferred to a 96-well plate (Microtest Plate 96-Well, Sarstedt) and incubated at 30°C for 24 h, 48 h and 72h. After each time point, the supernatant was carefully removed and cells were washed thrice with 1x PBS. 100 µl 0.1% crystal violet were added and the plates incubated at 37°C for 30 min. The supernatant was removed and cells were washed four times with 1x PBS. 100 µl 95% ethanol were added and the plates incubated at room temperature for 30 min at 50 rpm. Finally, the OD_595_ was measured using a BioTek Epoch 2 microplate reader. Biological triplicates were prepared with six technical replicates each.

### Growth assays

Growth assays were conducted as previously described (17). Briefly, overnight cultures were used to inoculate 10 ml of BHI medium to an initial OD_600_ of 0.1. Cultures were incubated at 37°C at 200 rpm until reaching an OD_600_ of 0.3. 2 ml of each culture per strain and test condition were harvested by centrifugation. The cell pellets were washed twice with LSM medium containing the respective sugars. Cultures were then adjusted to an OD_600_ of 0.1 and 200 µl of each transferred into a 96-well microplate. The plate was incubated at 37°C with orbital shaking in a BioTek Epoch 2 microplate reader and OD_600_ measurements were recorded every 15 min for at least 25 hours. Data from three independent experiments were averaged and mean values with standard deviations were plotted.

### Fluorescence and differential interference contrast (DIC) microscopy

For microscopy, *L. monocytogenes* strains EGD-e *treB*::*P_p60_-msfGFP* and EGD-e *treB*::*msfGFP* were grown overnight in 4 ml BHI at 37°C with agitation. Rhamnose-dependent promoter induction was assessed by culturing EGD-e *treB::P_rha_-msfGFP* strains in LB medium as *P_rha_* is subject to carbon catabolite repression (CCR) (13). Cultures were grown for 20 h at 37°C and 200 rpm in 4 ml LB medium supplemented with either no rhamnose or with rhamnose at final concentrations of 0.05%, 0.1%, 0.2% and 0.5% (v/v) as well as 0.5% rhamnose combined with 0.5% glucose. For microscopy, 1 µl of each culture was spotted on microscope slides covered with a thin agarose film (1.5% agarose in distilled water), air-dried and covered with a coverslip. Differential interference contrast (DIC) and fluorescence images were taken at a 100x magnification using an AxioImager M1 microscope (Zeiss, Jena, Germany) and images were captured using a Photometrix CoolSNAP HQ camera (Roper Scientific, Photometrics, Tucson, USA) and processed using ZEISS ZEN Digital Imaging (Version 2.3; Zeiss, Jena, Germany).

## Acknowledgements

We would like to thank Prof. Henrik Strahl (Newcastle University) for sharing plasmid pHJS105. We are grateful to Prof. Jörg Stülke for providing JR and JS with laboratory space, equipment and consumables and to the Göttingen Center for Molecular Biosciences (GZMB) for financial support. This work was funded by the German research Foundation (DFG) grant RI 2920/3-3 to JR.

